# Efficient and Robust Genomic DNA Isolation and Next-Generation Sequencing Library Preparation from Recalcitrant Wild Grape Species

**DOI:** 10.64898/2026.05.19.713680

**Authors:** Anugya Bhattarai, Jacob Smith, Heba Abdelgaffar, Rachel Carpenter, Swati Mishra, Juan Luis Jurat-Fuentes, Gautam Shirsekar

## Abstract

This protocol details the extraction of high-molecular-weight genomic DNA from grapevine tissues (wild and cultivated *Vitis* spp., including pathogen-infected samples) and the subsequent preparation of Illumina(R) whole-genome sequencing libraries using bead-bound Tn5 transposase. It is designed to overcome challenges from polyphenolic compounds and secondary metabolites in wild plants, providing a cost-effective workflow for large-scale population genomics. It includes recipes for buffers, incubation times, critical notes, and troubleshooting tips to maximize yield and library quality. Although designed for the grapevine DNA, this protocol is potentially applicable to other similar wild plant species

**Highlights:**

1. Optimized CTAB-PTB DNA extraction protocol for field-collected wild plant tissues.
2. Effective removal of polyphenols and secondary metabolites associated with DNA using PTB.
3. Cost-effective Illumina^Ⓡ^ DNA Prep library preparation using bead-bound Tn5 transposase (Tagmentation).
4. Scalable workflow suitable for large-scale population genomics in *Vitis* species.
5. Validated method for high-molecular-weight DNA and high-quality sequencing data.

**Graphical Abstract:** 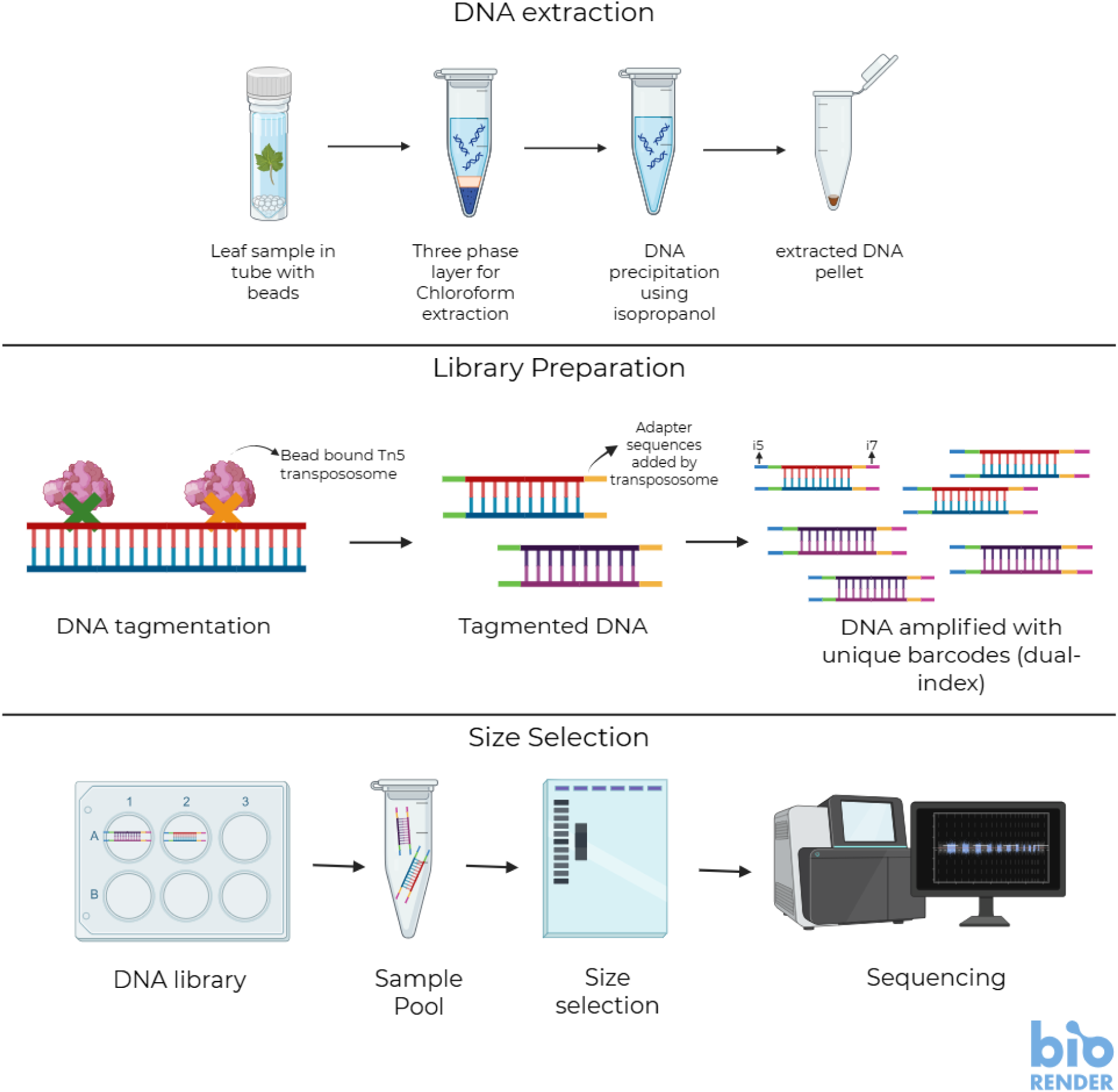

## Introduction

One way to study a biological system is to understand its genetic architecture. Use of genomics tools helps provide a comprehensive view of heritable variation, enabling researchers to identify causal variants underlying phenotypic traits, construct demographic histories, understand evolutionary patterns, and predict the system’s adaptive potential to environmental stimuli^1^. But what precedes the genomics inference is the information gathering step, i.e., DNA extraction, library preparation, and sequencing, which largely determines the quality of information obtained for these analyses^2^. Poor DNA quality leads to biased variant calls, reduced genome coverage, and ultimately compromised biological conclusions^2,3^.

For studies requiring large sample sets, e.g., population genetics studies, the sample is directly collected from the field without multiplying it in a controlled greenhouse setting^4^. The comprehensive sampling for this study needs coverage over a wide range of environmental gradients and geographic ranges, which makes controlled environment multiplications not very feasible^5^. However, unlike the greenhouse-grown or controlled-environment samples, field-collected plant tissues experience various biotic and abiotic stresses. These stresses can elevate levels of polyphenolic compounds, polysaccharides, and secondary metabolites that interfere with DNA extraction and downstream enzymatic reactions^6,7^. These contaminants co-precipitate with DNA during purification, inhibit polymerases during library preparation and PCR amplification, and reduce overall sequencing success rates^8^. This ultimately gives a poorer DNA library and more erroneous sequencing outcomes, which make inferences inaccurate and difficult.

The commercial extraction kits are mostly optimized for greenhouse-grown plant tissues and do not work well for wild samples^9^. Similarly, the DNA library prep kits are also optimized for specific DNA quality and are largely inflexible. Adding to it, the commercial DNA extraction kits are expensive, and per-sample library prep costs are prohibitive, especially for studies requiring a large number of samples^10^. Additionally, the need for extensive DNA quantity normalization, re-extraction of failed samples, and repeated library preparation attempts because of low-quality inputs further inflates costs and extends timelines ^11^.

To address these challenges, an integrated workflow is required that pairs a robust DNA extraction method for field-collected tissues with a whole-genome sequencing (WGS) library preparation protocol that tolerates variations in DNA quality and quantity. By eliminating the need for extensive normalization and cleanup, the workflow significantly reduces per-sample costs, making population-scale sampling feasible.

Therefore, we developed and optimized such a workflow using wild North American grape species (*Vitis* spp.) as our model system. Grape leaves are notoriously recalcitrant for DNA extraction due to exceptionally high concentrations of polyphenolic compounds (particularly in wild species adapted to high-light, water-stressed environments) and complex polysaccharide matrices^12^. If a DNA extraction protocol succeeds with field-collected grape tissue, it is likely to be broadly applicable to other challenging plant systems.

We combined a modified CTAB extraction^13^ with N-phenacylthiazolium bromide (PTB) for maximum DNA recovery from field-collected grapevine tissue. To pair with it, we have optimized the Illumina DNAprep protocol for DNA library preparation, which is a cost-effective and a DNA quantity/quality flexible option.

The CTAB-PTB extraction method builds on established plant DNA isolation chemistry but incorporates PTB; previously used primarily for ancient DNA and herbarium specimens to cleave glucose-derived protein crosslinks and release DNA trapped in polysaccharide-protein complexes that accumulate in stressed field tissues^14^. This modification consistently improves both DNA yield and purity (as measured by 260/230 absorbance ratios) from recalcitrant samples without requiring additional purification steps.

For library preparation, we employ a bead-bound Tn5 transposase approach that significantly reduces per-sample costs compared to commercial kits while maintaining compatibility with standard Illumina sequencing platforms. Critically, the tagmentation-based workflow tolerates DNA concentration variation across a broad range (1-60 ng/μL in our validation) and accommodates moderate quality variation (260/230 ratios of 1.2-2.2) without requiring extensive normalization or cleanup, making it well-suited to the heterogeneity inherent in field-collected samples. The bead-bound format further improves enzyme stability and enables efficient washing steps that remove contaminants, reducing library preparation failures^15^.

The complete workflow from field-collected leaf tissue to sequencing-ready libraries can be completed in 2.5 days with per-sample costs of approximately $3. We have consistently used this protocol in the lab for getting our sequenced results, which have worked for more than 500 samples now. The data range for 8 different samples has been provided in this protocol.

Although developed specifically for wild grapevine, this protocol is broadly applicable to other woody plant species with high polyphenolic content, particularly those sampled from natural populations where environmental stress is unavoidable. By enabling cost-effective, high-quality sequencing of large samples from field-collected material, this workflow removes a critical barrier to population-scale genomic studies in plant systems.

When to use this protocol:

– When working with field-collected plant tissues that are exposed to environmental stressors, resulting in the accumulation of secondary metabolites.
– The protocol (library prep) is optimized for plate format. Therefore, this protocol is useful for large sample sets (>50 samples) where cost reduction is important.

Permissions:

– The samples were collected from vineyards and the National Forest of Tennessee, with permission from vineyard owners and the forest service division. The permit for sampling across the National Forest can be provided upon request.

### DNA extraction (Day 1)

1. Prepare CTAB-PTB Buffer (50 mL Total, sufficient for 60 samples with 750ul of buffer/ sample + 5ml extra). Recipe in Table 1. Note: CTAB buffer without the PTB and β-mercaptoethanol can be prepared beforehand and stored for large samples at room temperature.
2. Aliquot the CTAB-PTB buffer based on the number of samples needed. Add the following chemicals immediately prior to use:

- β-mercaptoethanol (0.2%): Add 1.5 ul per sample
- PTB (0.063%): Add 0.47 mg per sample

#### 1. Grind Samples (10 minutes)

A new growth fully opened leaf tissue of about 50mg weight should be taken and kept in bead beater screw cap tubes with BioSpec Products 2.3 mm Zirconia/Silica Beads inside them. The sample is stored at −80 °C overnight or until you are ready to extract the DNA. (This step should be done beforehand to allow proper freezing of the tissue.) Once ready for grinding of samples stored in tubes, bring them out of the freezer and put them in Fisherbrand^TM^ Bead Mill 24 Homogenizer with the following settings:

- Speed (S): 6.00
- Time (T): 0:20s
- Cycles (C): 01
- Delay (D): 0:00

**Table 1.** CTAB buffer recipe.

|  | Final Concentration | Stock Concentration | For 50 ml | Remarks |
| --- | --- | --- | --- | --- |
| CTAB | 2% | - | 1 g |  |
| EDTA | 20mM | 0.5 M | 2 ml |  |
| Tris-Cl (pH 8.0) | 100mM | 1 M | 5 ml |  |
| NaCl | 1.4M | 5M | 14 ml |  |
| PVP | 0.64% | - | 320 mg |  |
| ddH<sub>2</sub>O |  | - | To 50 ml |  |

Note: *Take the samples out only when all preparations have been done, the CTAB buffer with PTB and β-mercaptoethanol has been made, and when you are ready to grind the samples to avoid thawing of tissues. Do not use liquid nitrogen on tubes; the labels will fall off (unless cryo-labels are used)*.

#### 2. Add CTAB-PTB Buffer (10 minutes)

- After the sample has been ground, immediately add <u>750 uL</u> of your aliquoted CTAB-PTB buffer to each tube.
- Mix thoroughly to make a uniform homogenate of the buffer and the ground tissue.

#### 3. Incubations (1 hr)

- At room temperature, rotate the tubes using the Fisherbrand™ Mini Tube Rotator for 30 min at a speed of 25 rpm.
- Immediately after rotations, incubate the tubes at 65 °C for 30 min.

#### 4. Chloroform Extraction (30 minutes)

- Take the sample to the fume hood and add 750 uL of 24:1 chloroform: isoamyl alcohol and mix by inverting the tubes gently 2-3 times.
- Centrifuge at 13,000 rpm (15,890 xg) for 15 min.
- Carefully transfer 500 uL of the top aqueous layer to a new tube.

Note: *Avoid the lower layer (insoluble phase and debris); contamination here will affect the downstream extraction and lower the quality and quantity of the DNA*.

#### 5. Isopropanol Precipitation (30 minutes)

- Add 500 uL ice-cold Isopropanol to the new tube. *(Isopropanol should be chilled for 15 minutes on ice before use.)*
- Gently invert 10–15 times until the precipitated DNA strands become visible.
- Centrifuge at 13,000 rpm (15,890 xg) for 15 min to pellet DNA.

#### 6. Ethanol Washes (Return to Bench) (45 minutes)

- Pour off supernatant, leaving the pellet.
- Add 500–1,000 uL of 70% freshly prepared ethanol.
- Dislodge the pellet by flicking the tube gently so that it floats freely.
- Centrifuge at 13,000 rpm(16,060 xg) for 5 min.
- Repeat the second Ethanol wash (500–1,000 uL 70% freshly prepared ethanol). Again, dislodge the pellet.
- Pour off Ethanol.
- Pipette out any leftover Ethanol completely.
- Air-dry with the lid open for ∼2 min.

#### 7. Elution (10 minutes)

Prepare elution mix:

- 50 uL elution buffer per sample(should be prepared before you start the protocol)
- Add 2 uL RNase A per sample to the elution buffer. (Final volume: 52 uL; add 51 uL to each pellet)
- Add 51 uL of the elution mix (Elution buffer + RNase A) to the DNA pellet.
- Let the DNA resuspend/relax in the buffer completely (lightly flick if needed) overnight at ambient room temperature.

**|| Pause:** pause for the day and carry over next day.

DNA Quantification (day 2) (∼1:30 hr)

Before measuring on Nanodrop or Qubit:

- Incubate the samples at 37 °C for 1 hour.

Then proceed with the measurement of your DNA quality (on Nanodrop) and quantity (on Qubit).

**|| pause:** DNA can be stored in 4°C until you use it for library prep.

### A. Illumina Library Preparation

Protocol type: Wet lab

Scale: 1 × 96-well plate

Notes: *Calculations shown are for 1 plate unless otherwise stated*.

#### Tn5 preparation and transpososome assembly

We referred to the previously established protocol for Tn5 preparation^16^. Tn5 is also available commercially, but that increases the library prep cost. Purify the Tn5, prepare Tn5 duplexes(Tn5ME-A, Tn5ME-B, and Tn5ME-rev), and assemble the Tn5 transpososome according to the protocol^16^.

#### Overview

This protocol describes:

1. Preparation of bead-bound Tn5 transpososome (Step 0; stock prep)
2. Tagmentation reaction on beads (Step 1)
3. Stripping Tn5 from DNA (Step 2)
4. PCR amplification and indexing (Step 3)
5. Library QC by gel electrophoresis

#### REAGENTS USED

*Note: stocks can be prepared previously and stored, but all the buffers need to be prepared fresh on the day of the experiment*.

- Hydrophilic Streptavidin-binding beads (catalogue number: S1421S): 2.5ul/sample (stored at 4 °C, need to be taken out to room temperature before use)
- Assembled Tn5 transpososome: 0.1ul/sample (stored in −20 °C). The amount per sample to be used needs to be tested for each batch of Tn5 prepared.
- Dialysis/Storage Buffer:

*Note: DTT should be added fresh. Add in the order mentioned below in Table 4*.

- Streptavidin binding buffer
- Wash Buffer (WB)
- Wash buffer with SDS (WB-SDS):

### STEP 0: Binding Transpososome to Magnetic Beads (Stock Preparation) (2 hrs)

*Note: This step can be done in advance. The end product can be stored at 4°C for up to 1 month without significant loss of Tn5 activity*.

Calculations:

- Scale: 1000 samples
- Streptavidin Binding Beads (catalog number: S1421S): 2.5 ul/sample = 2500 ul total

#### 1. Preparing the beads

- Remove Streptavidin binding beads from fridge and keep at room temperature, 25 °C for 30 minutes
- Take 2.5 ul stock beads/samples: 2.5 * 1000 = 2500 ul beads in a 50 mL Falcon tube (stock beads need to be at RT, shake/vortex the bottle well before adding to the tube)
- Put the falcon tube in a magnetic stand and remove the storage buffer.
- Do not forget to return the stock bead bottle to the fridge

#### 2. Wash beads

- Add 4X volume of Streptavidin binding buffer to the tube. (For 2500ul of beads: 2500*4 = 10000ul of the buffer)
- Quick vortex to resuspend the bead and put it on the magnet to remove the buffer

#### 3. Add enzyme

- Add 6X volume of the Streptavidin binding buffer to the tube (for 1000 samples: 2500 * 6 = 15000 ul of the buffer)
- add the calculated amount of purified assembled Tn5 transpososome (0.1 ul per reaction i.e, 0.1 * 1000 = 100ul for 1000 samples)

#### 4. Incubate

- Close the tube, mix by inverting, and let it rotate at RT for 30 minutes at 10rpm on the Fisherbrand™ Mini Tube Rotator.
- Then, put on a magnetic rack for 2 minutes and remove the buffer.

#### 5. Change buffer

- Add 6X volume of dialysis buffer and rotate with the Fisherbrand™ Mini Tube Rotator at 10 rpm at RT for 5 minutes. (for 1000 samples, 2500 * 6 = 15000 ul)
- Put it on the magnet for 2 minutes and remove the buffer

#### 6. Resuspend and store

- Resuspend the beads in 4X volume of Dialysis buffer (for 1000 samples: 2500*4 = 10000ul)
- For storage: for easier resuspension, store the Bead-Bound Tn5 in 2ml Eppendorf tubes; avoid storing in the Falcon tubes. store at 4 °C)

**|| pause:** can be paused here and resumed the next day

### STEP 1: Tagmentation Reaction with Bead-Bound Tn5 (1 hr)

Before Starting

- Set the thermocycler to 55 °C for 10 min
- Keep the 96-well magnetic rack on ice
- Prepare

- TAPS-DMF-MgCl₂,
- wash buffer (WB)
- wash buffer + SDS (WB-SDS)
- Thaw i5, i7 primers, and dNTPs before use.
- Keep MixMate at room temperature (25 °C)

**1.** Prepare Beads for 96 Samples

- Transfer 1200 ul bead stock (sufficient for ∼120 samples) into a 2 ml tube.
- Place on the magnet for 2 min, remove supernatant.
- Add 600 ul wash buffer and resuspend.
- Aliquot ∼72–75 ul into PCR strip tubes.
- Keep PCR strip tubes on ice.
**2.** DNA Transfer and Tagmentation Reaction Setup

- Place a new 96-well plate on ice.
- Centrifuge the DNA plate briefly.
- Add to each well:

- 3 ul DNA (The lowest concentration of DNA we could generate the library from was 1.07 ng/ul, the highest 60 ng/ul, measured on Qubit. We recommend using fluorophore-based DNA quantification for accurate estimation, but for a cheaper alternative, nanodrops can also be used. With quality ratio range: A260/280: 1.5-2.0, A260/230: 1.00-2.25, Lowest Concentration: 50ng/ul)
- 2 ul TAPS-DMF-MgCl₂
- Centrifuge the plate with DNA and TAPS-DMF-MgCl₂ briefly for 5 seconds
- Add 5 ul beads to each well (<u>work quickly on ice</u>).
- Ensure beads contact DNA (centrifuge if needed).
**3.** Mixing & Incubation

- Cover the plate with a PCR cover(Thermo Scientific™ Adhesive PCR Plate Foils) (on ice).
- Briefly place the plate on a 96-well magnetic rack (∼5 seconds)
- Mix on Eppendorf™ ThermoMixer Temperature Control Device (1500–2000 rpm) for ∼30 seconds (check if it is mixed properly, increase the time by 15 seconds if not mixed well)
- Incubate at 55 °C for 10 min in the PCR machine.

### STEP 2: Stripping Transpososome from DNA (30 minutes)

#### 1. SDS-mediated denaturation

*Do NOT perform this step on ice, as SDS precipitates on ice*.

- Add 10 ul wash buffer + SDS (WB-SDS) to each well.
- Mix on the Eppendorf™ ThermoMixer Temperature Control Device (∼15 seconds, 1500-2000rpm).
- Incubate at 55 °C for 5 min in the PCR machine.

#### 2. Washes

- Place the plate on a 96-well magnetic rack and remove the supernatant.
- Add a 150 ul wash buffer (no SDS). (WB)
- Incubate ∼1 min at RT.
- Place on a 96-well magnetic rack, wait until the beads attach to the side walls, and remove the wash buffer.
- Add a 100 ul wash buffer (no SDS). (WB)
- Hold until the PCR setup is ready.
- Before PCR, place the plate on a 96-well magnetic rack and remove the wash buffer <u>completely</u>. (If there is some leftover buffer, use 10 ul tips to remove the buffer completely)

### STEP 3: PCR Amplification & Indexing (1.5 hrs)

Take a 5ml Eppendorf tube and prepare the master mix as mentioned in Table 8.

*Note: all PCR reagents preparation should be prepared on ice*.

### Mastermix preparation

Indexing

#### 1. i5 (8 different indexes)

- Distribute the 220 ul of mastermix prepared in each of the wells of the 8-well PCR strip tube (250 ul capacity).
- Add 9.75ul of eight different i5 indexes into the individual wells of the 8-well PCR strip tube with the mastermix.

#### 2. i7 (12 different indexes)

- Remove the wash buffer from the 96-well plate containing tagmented DNA (step 2.2)
- Add 7.5 ul of i7 oligos to each well.

– In this particular setup for 96 samples, we will use the same i7 index along the rows and a different i7 index across columns. This strategy, coupled with unique i5 indexes in rows (step 3.1), results in 96 unique i5/i7 combinations for demultiplexing.
- Transfer 17.5 ul i5-PCR mix (from 8-well PCR strip tubes) to each well of the corresponding row of the plate above. (rowA of the 8-well PCR strip tube to rowA of the 96-well plate and so on)
- Mix by pipetting twice.
- Briefly centrifuge if needed.
- Cover the 96-well plate with PCR cover (Thermo Scientific™ Adhesive PCR Plate Foils) and place briefly on a 96-well magnetic rack (∼5 seconds).
- Mix thoroughly on Eppendorf™ ThermoMixer Temperature Control Device (30 seconds, 1500-2000 rpm)
- Place in a PCR machine with heated lids.

PCR cycle:

### STEP 4: Library Quality Check using Agarose Gel (1.5 hrs)

- Gel: 1% agarose
- Stain: SYBR Safe (6 ul per 100 mL gel)
- Sample:

- 5 ul library
- 1 ul loading dye
- Ladder: 1 kb ladder (100 bp increments)[Thermo Scientific™ GeneRuler DNA Ladder Mix]
- Run the gel at 150 V for 45 minutes and observe in the gel doc.
- Check for similarity in fragment distribution and intensity. Based on band intensity, adjust the later pooling volume accordingly.

**|| pause:** can be paused and resumed the next day or when all libraries are ready for pooling.

### C. Pooling and Two-Sided SPRI Size Selection of Indexed Libraries

#### Materials

- PCR-amplified indexed libraries
- SPRI beads, homemade (AMPure XP or SPRIselect can be commercially bought)
- Freshly prepared 80% ethanol
- Invitrogen™ Nuclease-Free Water
- 1.5 mL Eppendorf Low-bind tubes
- 96-well plate magnetic rack
- 2ml tube magnetic rack
- 1000 ul, 200 ul, and 10 ul pipettes and filtered tips
- Agarose gel electrophoresis setup (for QC)

#### Critical Notes

- Always use low-bind tubes to maximize DNA recovery.
- Bring SPRI beads to room temperature for ∼30 minutes before use.
- Pipette slowly and precisely to avoid DNA shearing and ratio errors.
- Avoid disturbing bead pellets during supernatant removal.

### STEP 5: Pooling of Libraries

#### 1. Pooling Strategy (30 minutes)

- Briefly centrifuge the library plate.
- For each library, transfer 6 uL of tagmented - PCR amplified DNA library into the pooling tube. (2ml low-bind tube). If there is high/low intensity from step 4, adjust this volume accordingly.

*Note: Although Nanodrop concentrations may guide pooling, agarose gel visualization provides a more accurate estimate of relative library abundance*.

- Store leftover libraries at −20 °C as backup.

### STEP 6: Determination of SPRI Bead Ratios

The bead-to-sample ratios used in this protocol (0.5× and 0.75×) were empirically selected based on agarose gel analysis of size-selection test reactions for a specific batch of homemade SPRI beads. Multiple bead ratios were evaluated, and the resulting fragment distributions were compared to a validated reference library. The chosen ratio combination provided optimal enrichment of the desired fragment size range for Illumina short read sequencing while maximizing the exclusion of both large (>700 bp) and small (<300 bp) DNA fragments.

*Note: SPRI bead behavior depends on DNA concentration, fragment distribution, and experimental goals. Users seeking different insert size ranges should perform a similar gel-based calibration and adjust bead ratios accordingly*.

### STEP 7: First Size-Selection (0.5×: Removal of Large Fragments) (20 minutes)

At 0.5×, large(>700 bp) DNA fragments bind to the beads, and smaller fragments remain in the supernatant.

- Measure pooled library volume (X uL).
- Add 0.5 * X uL of SPRI beads to the pooled library.
- Mix by gently flicking the tube until the beads are evenly homogenized.
- Incubate at room temperature for 5 minutes.
- Place the tube on a 2 ml magnetic rack until the solution clears(the beads are bound to the wall of the 2 ml low-bind tube).
- Carefully transfer the supernatant to a new low-bind tube. (<u>Volume = Y ul</u>)
- Discard the beads (contains large fragments).

### STEP 8: Second Size-Selection Cut (0.75×: Capture of Desired Fragments) (20 minutes)

At **0.75×**, DNA fragments within the desired size range(>300 bp) bind to the beads, while smaller fragments(<300 bp) remain in solution.

- Add 0.25 * <u>Y ul</u> SPRI beads to the recovered supernatant. (Since 0.5 is already added before, this time just add 0.25X beads to make the overall bead concentration 0.75X)
- Mix gently and incubate at room temperature for 5 minutes.
- Place on a 2 ml magnetic rack
- Carefully remove and discard the supernatant.

- Beads now contain the desired DNA fragments(300-700 bp).

### STEP 9: Bead Washing

- Keep the tube on the 2 ml magnet.
- Add 80% ethanol without disturbing the beads.
- Incubate for ∼30 seconds, then remove the ethanol.
- Repeat for a total of two washes.
- Air-dry beads briefly for 2-5 minutes (do not over-dry).

- Note: If the beads begin to look flakey then they are being over-dried

### STEP 10: DNA Elution

- Remove the tube from the 2 ml magnet rack.
- Add 20–30 uL nuclease-free water.
- Resuspend the beads thoroughly by gently flicking the tube.
- Incubate at room temperature for 2–5 minutes to allow the DNA to resuspend in the water.
- Place the tube back on the 2 ml magnetic rack.
- Transfer supernatant (pooled ready-to-sequence libraries) to a new low-bind tube.

### STEP 11: Quality Control (1 hr)

- Analyze 1–2 uL of the final library on:

- 1% agarose gel or
- Bioanalyzer/TapeStation
- Compare fragment distribution to:

- DNA ladder
- Previously validated reference library (if available)

## Results

### 1. DNA yield and purity

As a validation for the protocol, we had 8 different wild and cultivated grape tissues(Table 11) whose DNA was extracted using our protocol. The Nanodrop concentrations ranged from 153.8 to 1074 ng/µL, while Qubit measurements ranged from 2.17 to 59 ng/µL, illustrating the broad range of DNA yield obtained from both greenhouse and field-collected grape tissues (Table 11). A260/280 ratios were generally within acceptable limits (1.5–1.97), indicating relatively low protein contamination across samples. In contrast, A260/230 ratios varied widely from 0.61 to 1.67, with several field-collected samples (e.g., Sus1, TP1, NC9) falling below the commonly recommended threshold of 2.0 for high-purity DNA, consistent with residual polysaccharides and polyphenolic contaminants in stressed wild tissues (Table 11). This range is not particularly preferred for the previously described DNA library preparation methods, which require a high quality and quantity of DNA^17^. But our described protocol, even with suboptimal 260/230 ratio, yielded sufficiently high-molecular-weight DNA for downstream library preparation and also produced a good library(Figure 6), indicating that the CTAB–PTB extraction is robust to moderate levels of co-extracted metabolites and the tagmentation-based protocol is flexible for suboptimal DNA quality.

### 2. Gel visualization

A 1% agarose gel visualization of the extracted DNA showed distinct, predominantly high-molecular-weight bands for all samples, with minimal smearing, consistent with limited mechanical shearing during extraction (Figure 5). The presence of strong bands in both greenhouse- and field-collected samples indicates that the CTAB–PTB protocol reliably recovers intact genomic DNA, even from recalcitrant wild *Vitis* tissues. (Table 11, Figure 5).

### 3. Library preparation and pre–size-selection profiles

Following tagmentation and PCR amplification, 1% agarose gel electrophoresis of the sequencing libraries revealed the expected smear characteristic of Illumina-compatible fragment distributions. Although the initial fragment range extended beyond the ideal window for short-read sequencing (Figure 6), these prepared libraries contain sufficiently amplified and within-range DNA fragments that can undergo successful size selection after pooling. The libraries from all eight DNA samples produced visible smears, confirming successful tagmentation and amplification despite the heterogeneity in input concentration and purity (Table 11, Figure 6). Notably, even samples with low A260/230 ratios (e.g., Sus1 and NC9) produced libraries with fragment distributions comparable to those from higher-purity greenhouse DNA, suggesting that the bead-bound Tn5 workflow effectively buffers against moderate secondary-metabolite carryover (Table 11, Figure 6).

### 4. Size selection and TapeStation quality control

Two-sided SPRI size selection (0.5× followed by 0.75×) consistently enriched fragments in the 300–700 bp range, as confirmed by both agarose gel and TapeStation analysis of the pooled library (Figure 7a–b). Post–size-selection gels showed the removal of high-molecular-weight fragments and reduction of low-molecular-weight noise, yielding a tight smear in the expected size range for Illumina short-read sequencing (Figure 7a). TapeStation electropherogram of the pooled library showed a primary peak centered at 496 bp, with minimal off-target peaks, and a final concentration of 6.545 ng/µL corresponding to 26.73 nM, indicating that the workflow reliably produces sequencing-ready libraries with homogeneous insert size distributions suitable for high-throughput population genomics (Figure 7b).

### 5. Sequencing Performance

Sequencing on the Illumina NovaSeq X platform generated between approximately 6.3 × 10⁷ and 1.1 × 10⁸ reads per sample, demonstrating high yield across all eight libraries (Table 12). Indexing performance was similarly robust: for all samples, roughly 95–96% of reads carried a perfect index sequence, with about 4–5% carrying a single index mismatch and negligible reads with two mismatches, ensuring accurate demultiplexing and minimal cross-contamination (Table 12). Importantly, there was no systematic reduction in read yield or index quality for samples with lower A260/230 ratios, indicating that the combined CTAB–PTB extraction and bead-bound Tn5 library preparation pipeline tolerates a wide range of input DNA purity while still producing high-quality sequencing data (Tables 11–12). Thus, even though several field-derived DNA samples did not meet conventional “ideal” purity criteria, they nonetheless generated high-quality libraries and sequencing outputs, underscoring that this protocol is specifically tailored to accommodate broad variation in DNA quality, particularly in A260/230 ratios (Tables 11–12, Figures 4–6).

**Figure 1:**
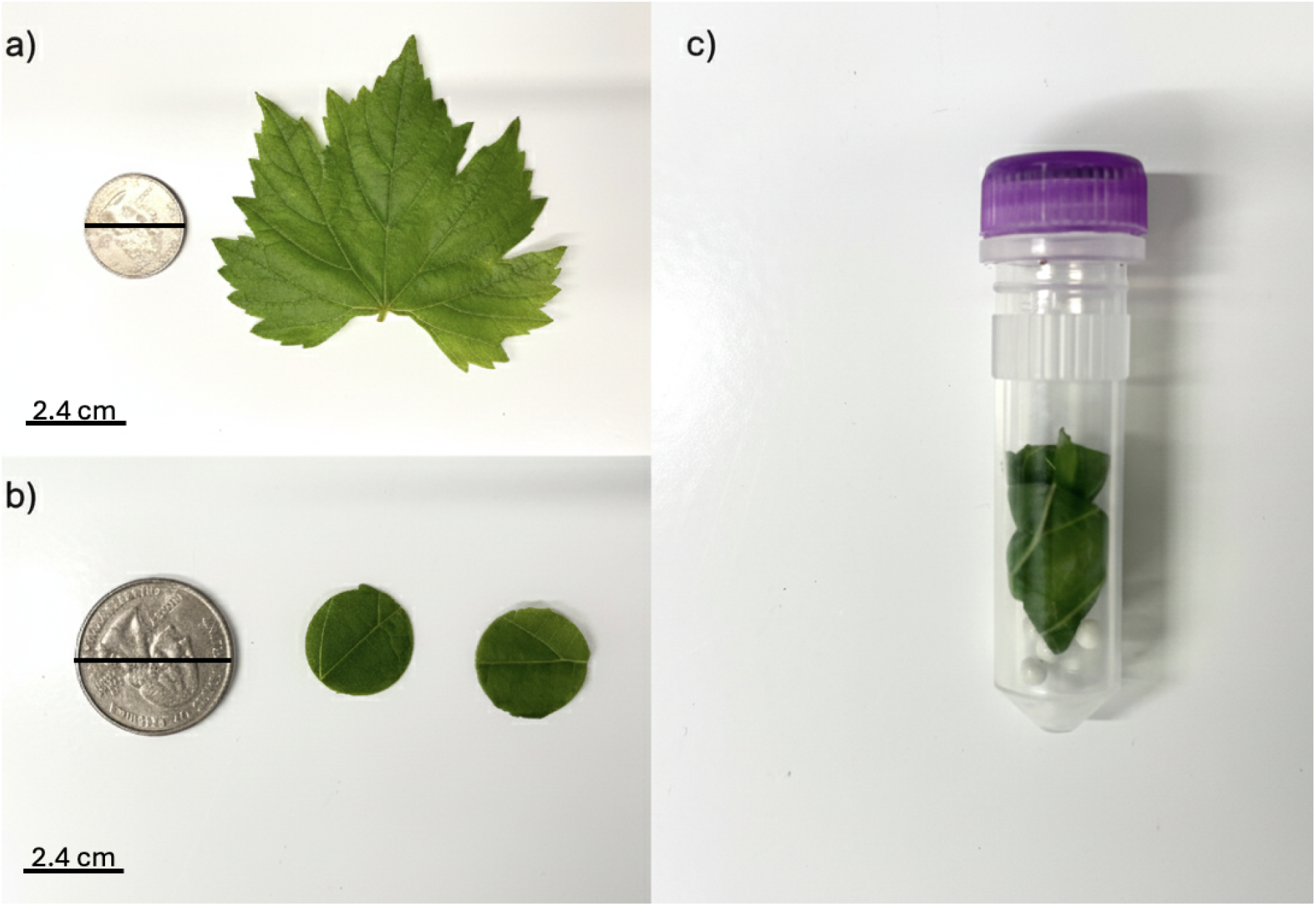
(a) Representative leaf used in this protocol with a 2.4 cm US Quarter as a size reference. (b) 1 cm Leaf disks of 1m diameter, cut from the leaf in Figure 1a, with a 2.4 cm US quarter as a size reference, for further DNA extraction. (c) Leaf disks from Figure 1b were transferred to the 2.0 bead beater screw cap tube for extraction.

**Figure 2.**
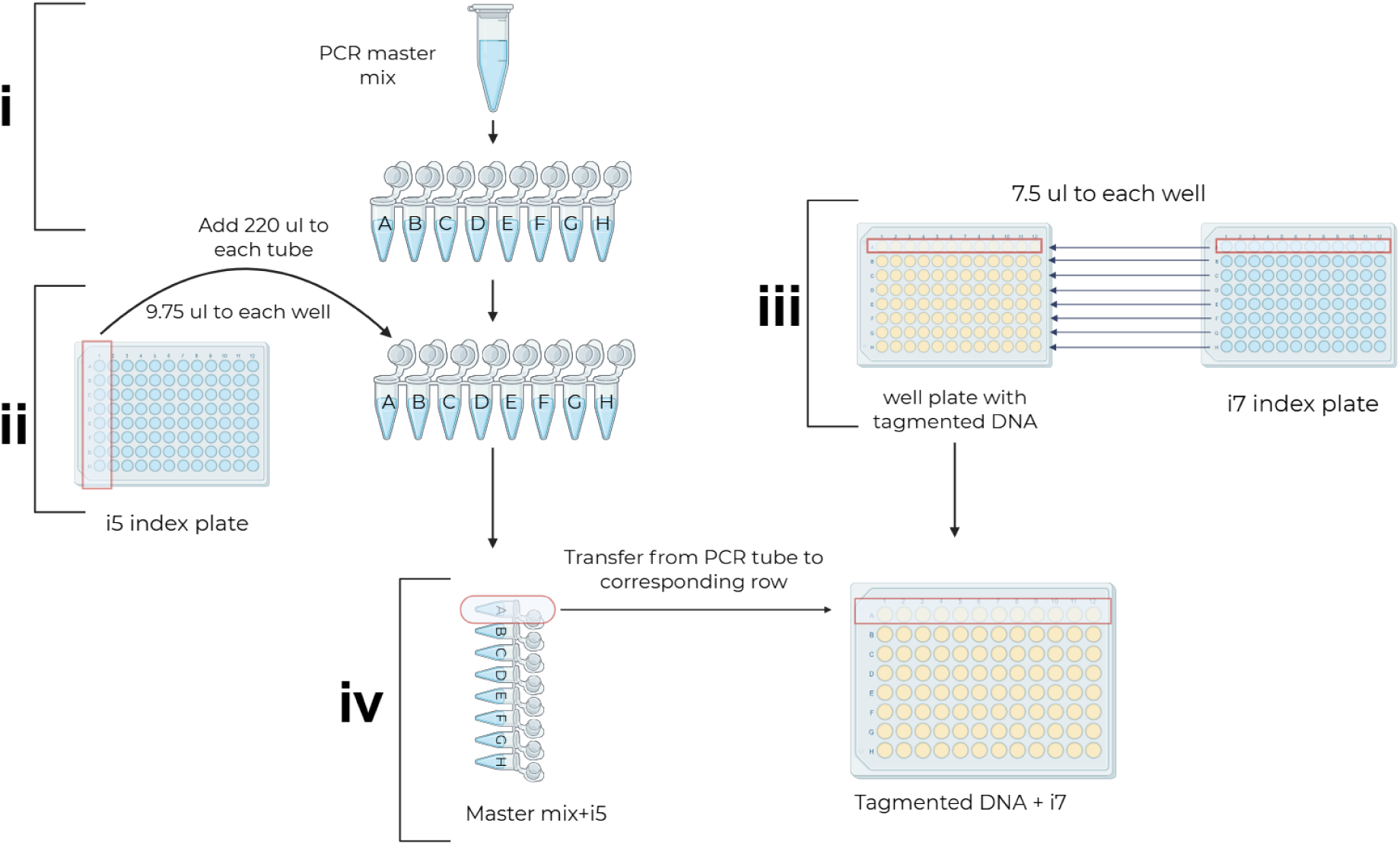
Indexing setup and PCR amplification of tagmented DNA. Step i. Preparation PCR mastermix and divide it into PCR strip tubes. Step ii. Addition of the i5 index to the mastermix. Step iii. Addition of the i7 index to tagmented DNA plate. Step iv. Addition of mastermix + i5 to the tagmented DNA plate that contains the i7 from step iii.

**Figure 3.**
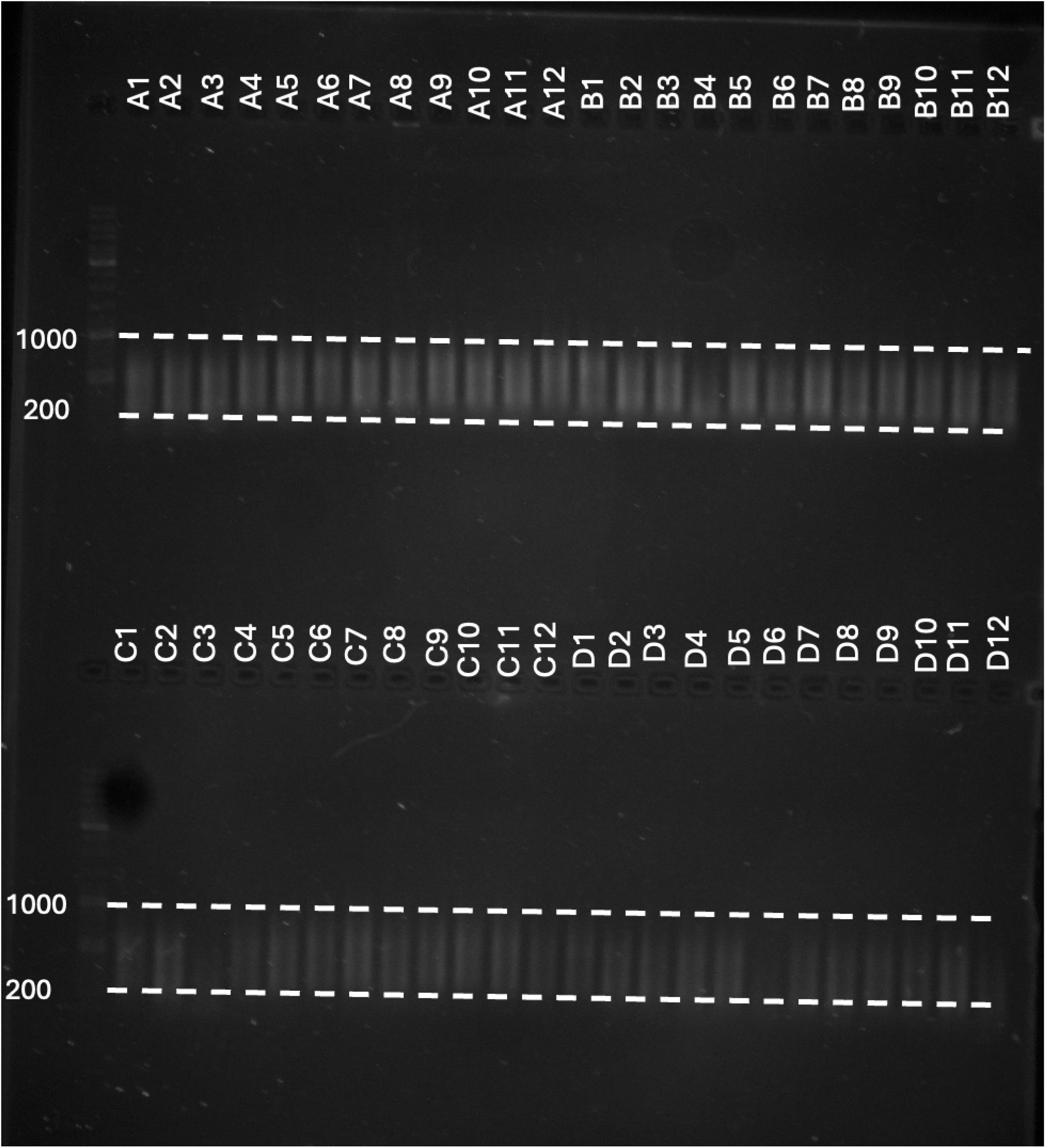
A 1% agarose gel electrophoresis of a successfully prepared DNA library run.

**Figure 4:**
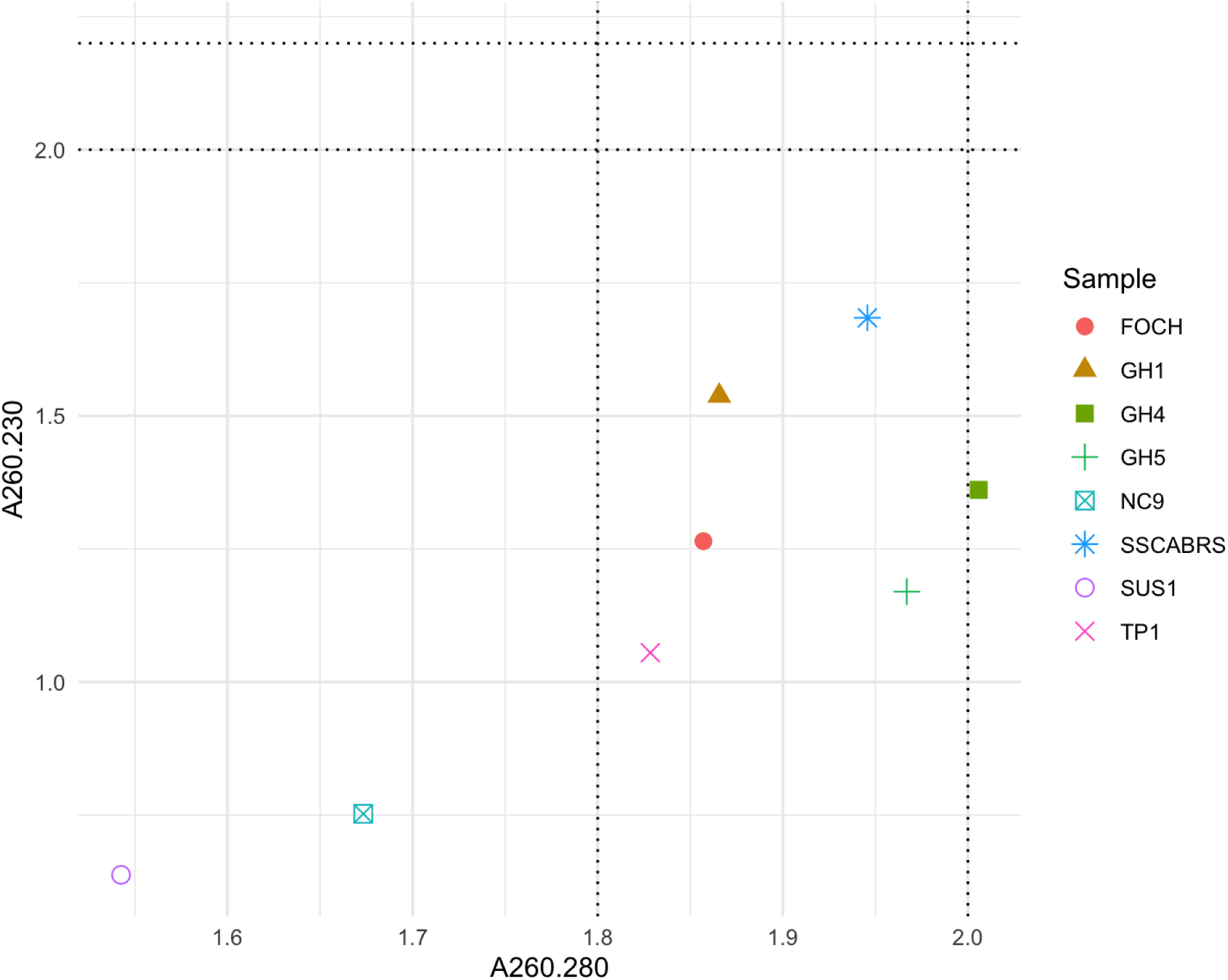
DNA quality was assessed using a NanoDrop spectrophotometer. Each point represents an individual sample, plotted by A260/280 (x-axis) and A260/230 (y-axis) ratios. The vertical dashed lines indicate the expected purity range for A260/280 (∼1.8–2.0), while the horizontal dashed lines indicate the expected purity range for A260/230 (∼2.0–2.2). Samples clustering within these ranges are considered high-quality DNA, whereas those falling outside may indicate contamination.

**Figure 5.**
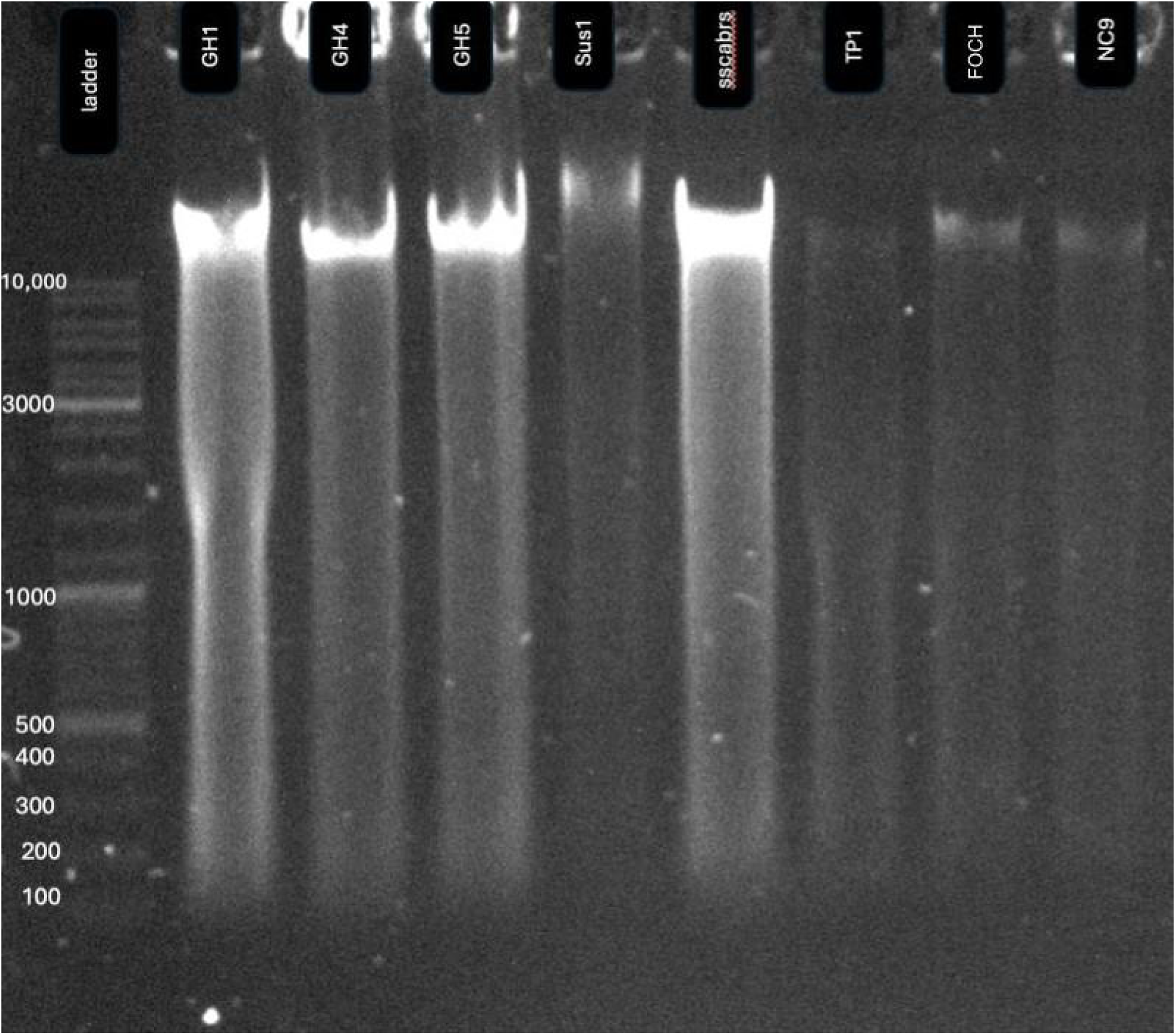
1% agarose gel electrophoresis of DNA samples run alongside a 1 kb DNA ladder. DNA bands are visible across the samples.

**Figure 6.**
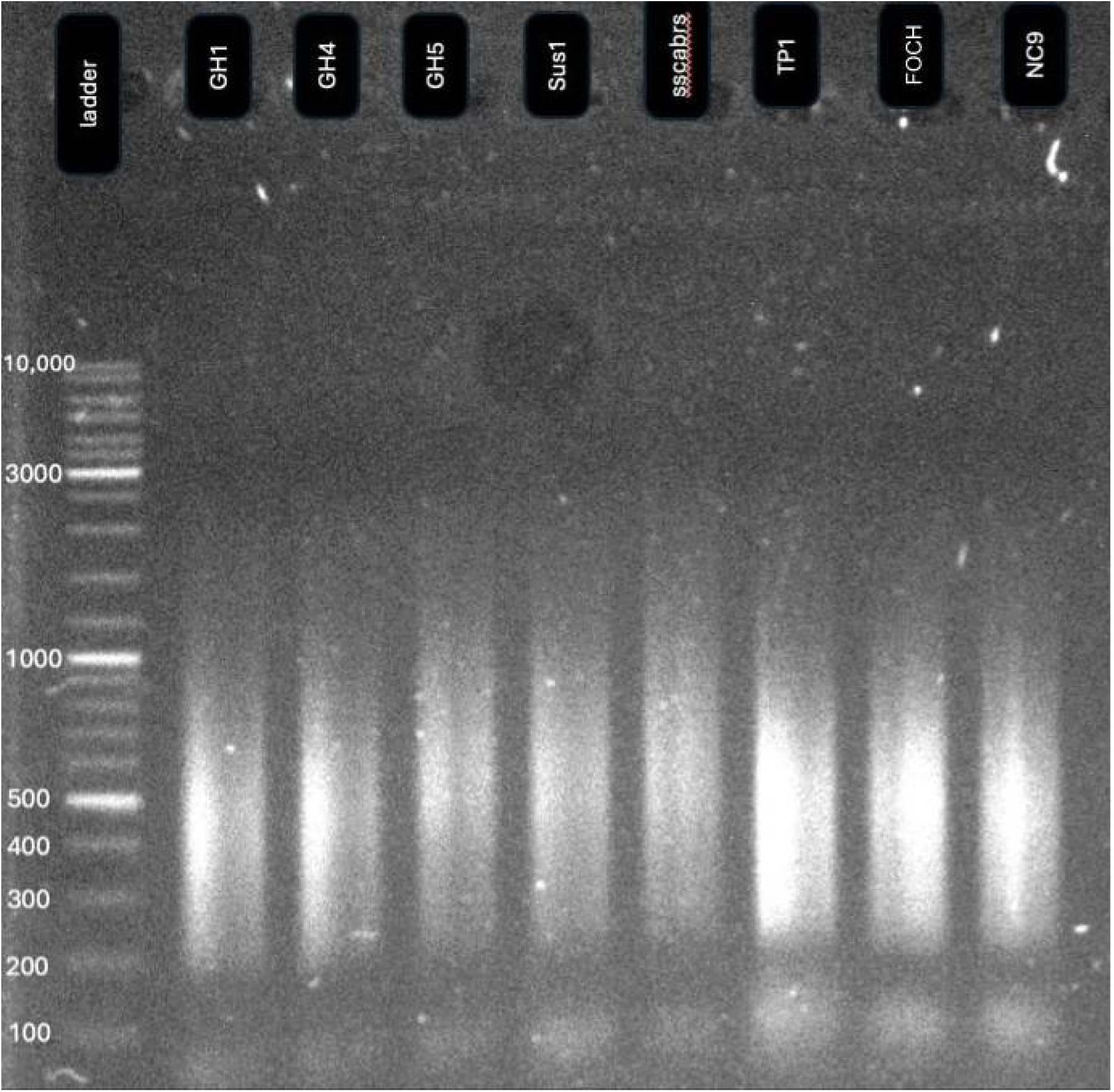
1% agarose gel electrophoresis of prepared sequencing libraries run alongside a 1 kb DNA ladder. The library shows a fragment distribution outside the desired size range for sequencing, which will be corrected through subsequent size selection steps.

**Figure 7.**
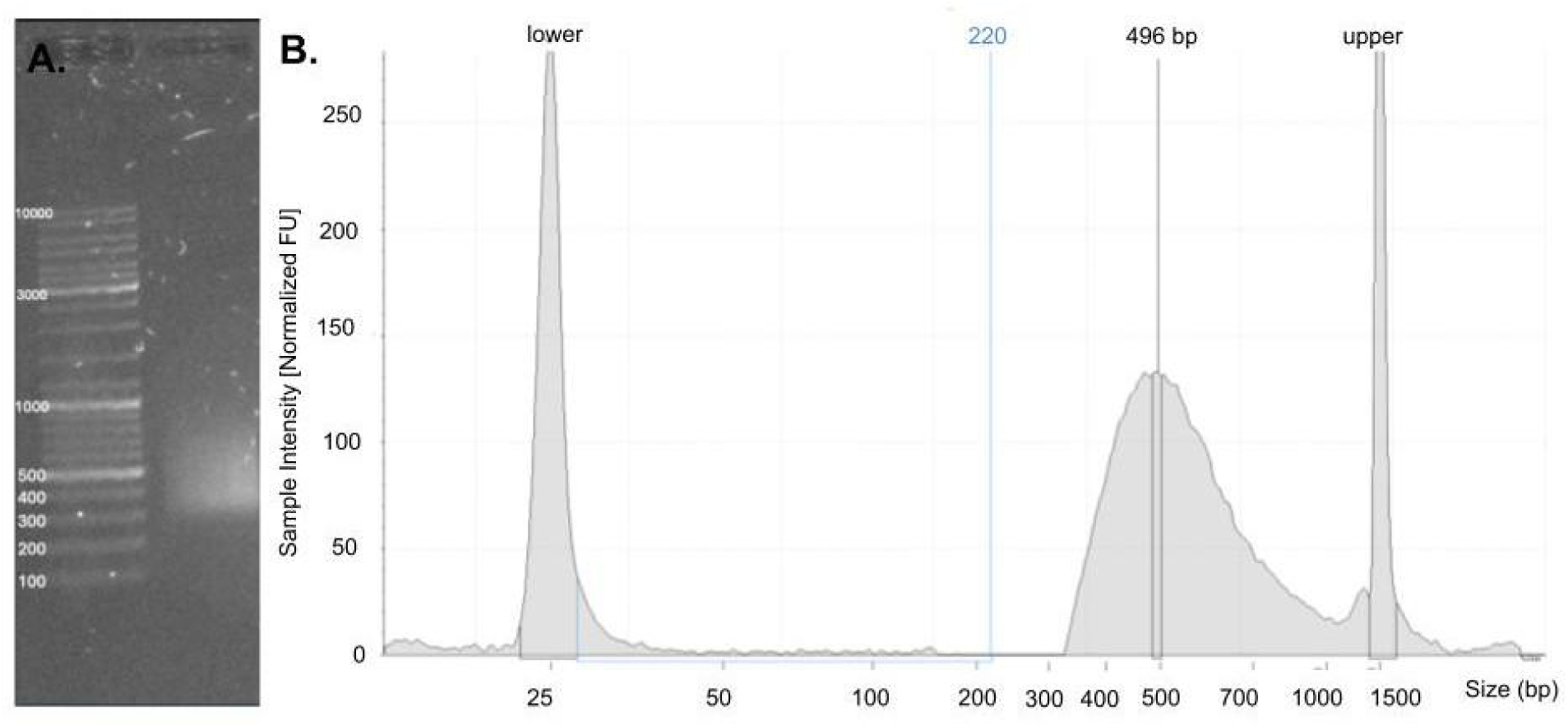
a) Successful pooled library size selection visualized on gel electrophoresis with a 1 kb ladder. b) A corresponding TapeStation analysis of the pooled library. The TapeStation electropherogram displays fragment size (bp) on the x-axis and normalized fluorescence intensity (FU) on the y-axis. The primary fragment peak is centered at 496 bp. The pooled library had a concentration of 6.545 ng/µL and a calculated molarity of 26.73 nM.

**Table 2.**
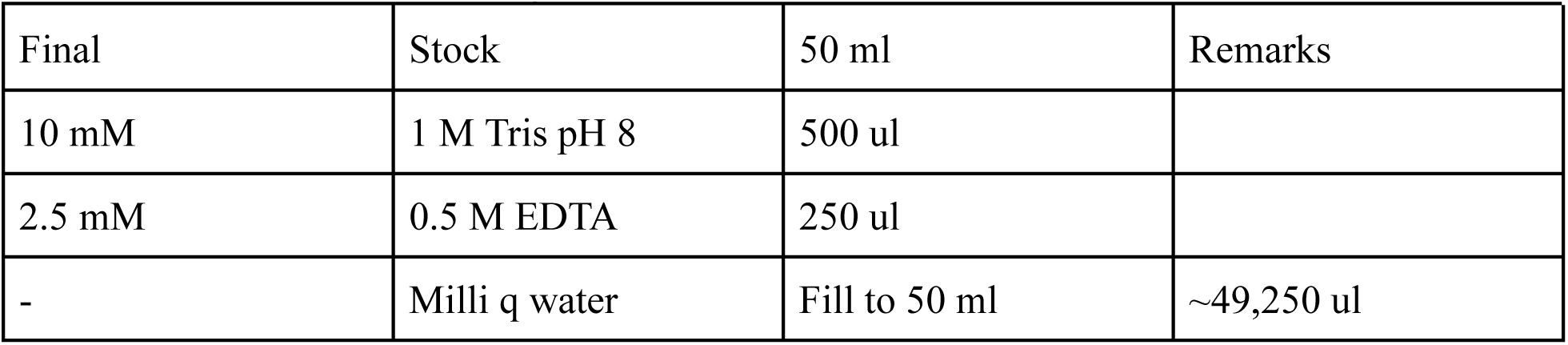
DNA elution buffer recipe.

**Table 3.** 5xTAPS-DMF-MgCl2 buffer recipe:

| Final | Stock | For 1ml | Remarks |
| --- | --- | --- | --- |
|  | Water (ddH<sub>2</sub>O, autoclaved) | 0.425 ml (425ul) | Mix in order listed here |
| 50mM TAPS | 1M TAPS; pH = 8.5(pH adjusted with 10M NaOH)<br>Sigma T5130,<br>243.3 g/mol | 50ul | water>TAPS>MgCl<br>2>DMF (gets warm, keep on ice) |
| 25mM MgCl<sub>2</sub> | 1M | 25ul |  |
| 50% DMF | 100% DMF | 500ul |  |

**Table 4.**
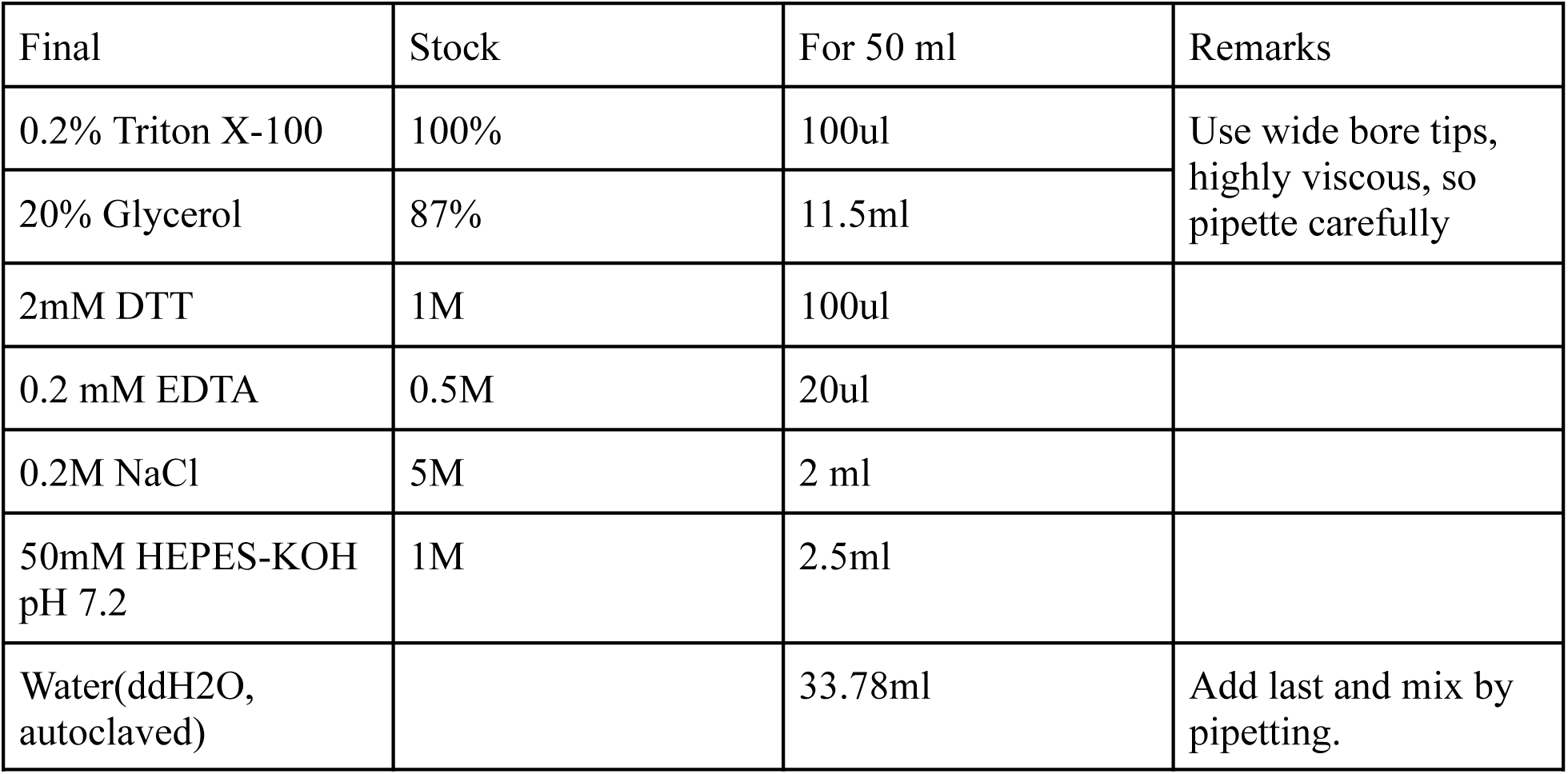
Dialysis/Storage buffer recipe.

**Table 5.** Streptavidin binding buffer recipe.

| Final | Stock | For 50 ml | Remarks |
| --- | --- | --- | --- |
| 0.1% Triton X-100 | 100% | 50ul | Viscous, use wide-bore tips |
| 0.5 mM EDTA | 0.5M | 50ul |  |
| 10 mM Tris, pH 8 | 1 M | 500ul |  |
| 0.6 M NaCl | 5M | 6ml |  |
| Water(ddH<sub>2</sub>O,<br>autoclaved) |  | 43.4ml |  |

**Table 6.** Wash Buffer recipe.

| Final | Stock | For 50ml | Remarks |
| --- | --- | --- | --- |
| 0.1% Triton X-100 |  | 50ul | Use wide-bore tips |
| 30mM Nacl | 5M | 300ul |  |
| 10mM Tris, pH8 | 1M | 500ul |  |
| Water(ddH<sub>2</sub>O,<br>autoclaved) |  | 49.15ml |  |

**Table 7.** Wash buffer with SDS recipe.

| Final | Stock | For 2ml |
| --- | --- | --- |
| 0.6% SDS | 10% | 120ul |
| Wash buffer |  | 1880ul |

**Table 8.** Mastermix recipe for tagmented DNA amplification.

|  | 1X | 112X (for 96 samples) | Remarks |
| --- | --- | --- | --- |
| **5X buffer** | 5ul | 560ul | Thaw it on ice immediately before use. |
| **dNTP** | 0.5ul | 56ul | Keep the thawed mix on the ice. |
| **Q5 polymerase** | 0.25ul | 28ul | take out from the freezer immediately before use. It is viscous, therefore pipette slowly and carefully |
| **Invitrogen™**<br>**Nuclease-Free Water** | 11ul | 1232ul | Add this last so that you can pipette and mix everything |
| **total** | 17.5ul in each tube |  |  |

**Table 9.** i5 index calculation to add to the mastermix.

|  | **1X** | **13X** | **Remarks** |
| --- | --- | --- | --- |
| **I5 index** | 0.75 ul | 9.75 ul | **(13X is for 12 columns of the 96-well plate + extra)** |

**Table 10.** PCR conditions for amplification of tagmented DNA.

| Temperature | Hold | Notes |
| --- | --- | --- |
| 72 °C | 10 min | Do not heat/denature tagmented DNA before this step |
| PAUSE |  | Pause the program, mix the plate in the Eppendorf™ ThermoMixer Temperature Control Device (1500-2000 rpm, 30 seconds, RT), then resume the PCR. |
| 98 °C | 30sec |  |
| 98 °C | 15sec |  |
| 65 °C | 20sec |  |
| 72 °C | 90sec | 11 cycles for the last 3 steps |
| end |  |  |

**Table 11.**
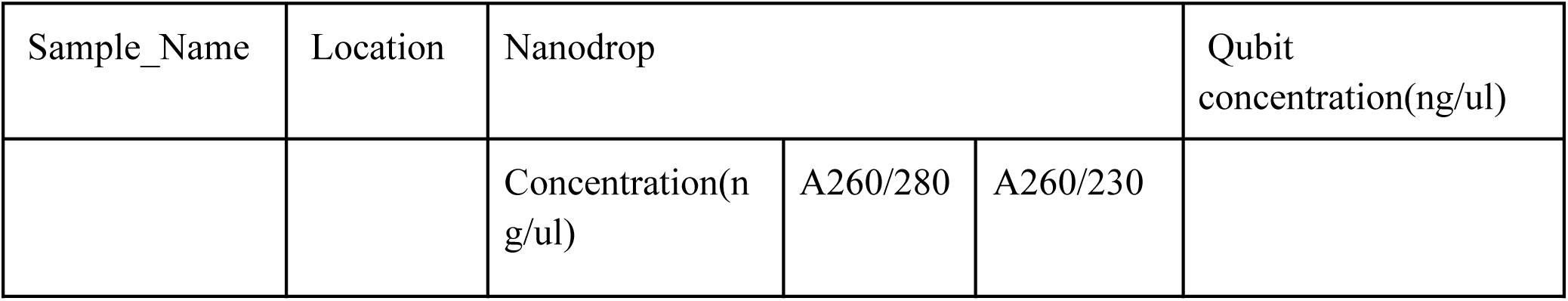
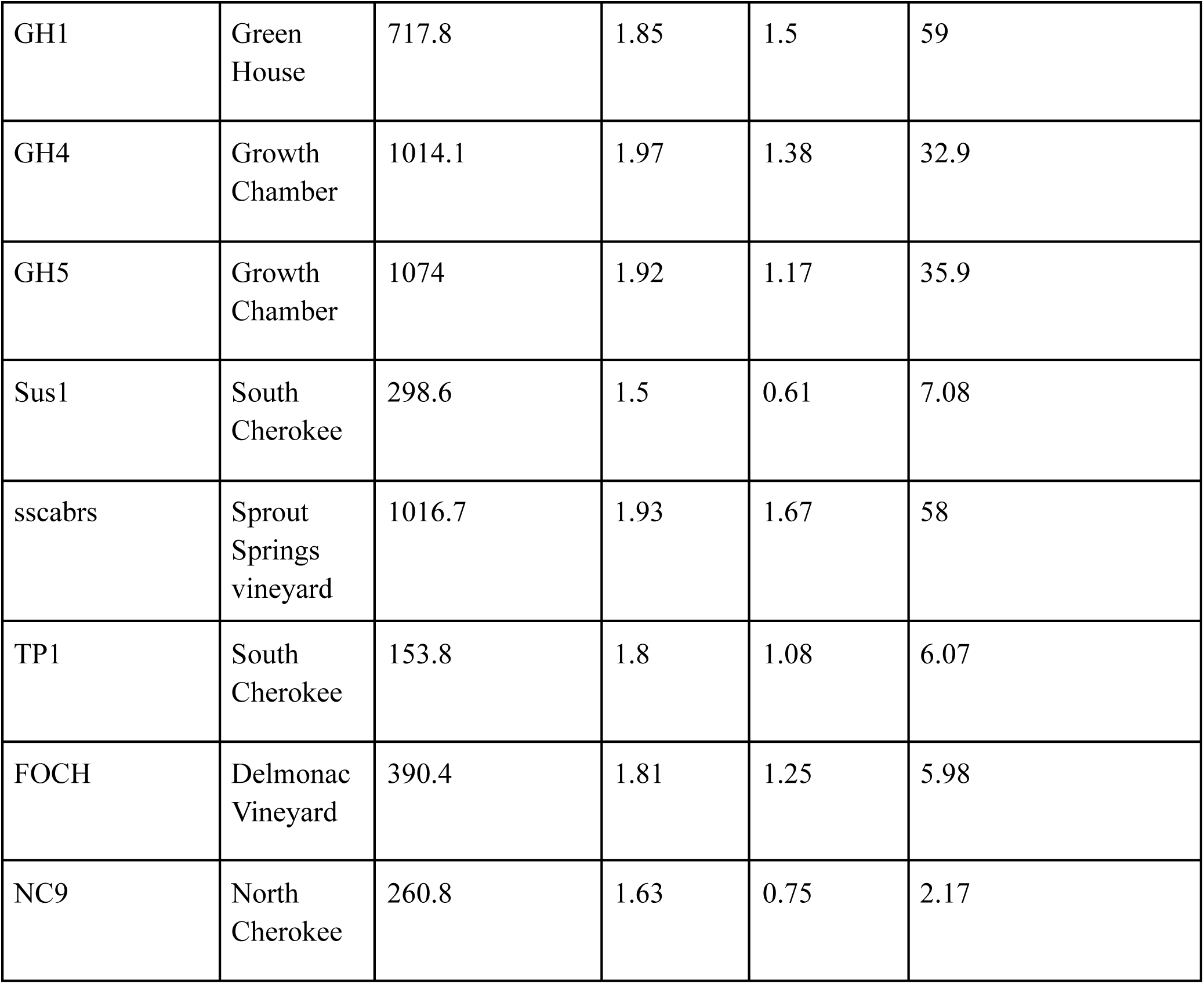
Qualitative and quantitative assessment of extracted DNA using Qubit and Nanodrop.

**Table 12.** Sequencing statistics from Illumina Novaseq X.

| Sample_<br>Name | TP1 | NC9 | SSCabrs | Sus1 | FOSH | GH1 | GH4 | GH5 |
| --- | --- | --- | --- | --- | --- | --- | --- | --- |
| Index | TATCAGCA<br>-TGAACCG<br>A | GATTC<br>GGA-TT<br>CTTCC<br>G | CACCT<br>ATC-GC<br>ATTGT<br>C | CCAAG<br>CAA-TT<br>CTTCC<br>G | CCAAG<br>CAA-G<br>CATTG<br>TC | ACGCT<br>AGA-G<br>CATTG<br>TC | ACGCT<br>AGA-AT<br>GCCTC<br>A | CTTGT<br>CTC-TG<br>AACCG<br>A |
| # Reads | 113551050 | 8321301<br>4 | 8249441<br>3 | 6275248<br>0 | 6478622<br>6 | 6857321<br>6 | 1102227<br>03 | 9818504<br>2 |
| # Perfect<br>Index<br>Reads | 108350549 | 7951728<br>0 | 7855381<br>8 | 5986094<br>8 | 6177516<br>7 | 6552502<br>4 | 1050498<br>92 | 9291125<br>7 |
| # One<br>Mismatch<br>Index<br>Reads | 5200501 | 3695734 | 3940595 | 2891532 | 3011059 | 3048192 | 5172811 | 5273785 |
| # Two<br>Mismatch<br>Index<br>Reads | 0 | 0 | 0 | 0 | 0 | 0 | 0 | 0 |
| % Reads | 0.0318 | 0.0233 | 0.0231 | 0.0176 | 0.0182 | 0.0192 | 0.0309 | 0.0275 |
| %<br>Perfect<br>Index<br>Reads | 0.9542 | 0.9556 | 0.9522 | 0.9539 | 0.9535 | 0.9555 | 0.9531 | 0.9463 |
| % One<br>Mismatch<br>Index<br>Reads | 0.0458 | 0.0444 | 0.0478 | 0.0461 | 0.0465 | 0.0445 | 0.0469 | 0.0537 |
| % Two<br>Mismatch<br>Index<br>Reads | 0 | 0 | 0 | 0 | 0 | 0 | 0 | 0 |

## Discussion

The integrated CTAB–PTB extraction and bead-linked Tn5 library preparation workflow provides a robust, low-cost route from field-collected wild *Vitis* tissue to sequencing-ready libraries, even when DNA purity deviates substantially from conventionally ideal metrics. By structuring the protocol sequentially from DNA extraction through library construction and two-sided SPRI size selection, we were able to maintain high-quality libraries while minimizing hands-on normalization and re-extraction steps.

High-quality DNA extraction and library construction are critical for successful whole-genome sequencing, yet these steps are often the most vulnerable to variation in sample quality, especially in field-collected plant tissues^2^. The field-collected plant tissues accumulate polyphenols, polysaccharides, and other secondary metabolites in response to various environmental stressors^8^. In our workflow, a modified CTAB backbone supplemented with N-phenacylthiazolium bromide (PTB) was sufficient to recover high-molecular-weight DNA across greenhouse and wild *Vitis* samples, despite wide variation in A260/230 ratios (approximately 0.6–1.7), indicating that the chemistry is tolerant of co-extracted metabolites that typically decrease purity scores. PTB likely contributes to this robustness by cleaving sugar-derived crosslinks and releasing DNA from polysaccharide-protein complexes in stressed tissues, reducing the extent to which polyphenols and polysaccharides remain tightly associated with genomic DNA and thus improving its accessibility to downstream enzymatic steps^14^.

The extracted DNA is converted into sequencing libraries using a bead-bound Tn5 transposase that both fragments the DNA and inserts adapter sequences in a single tagmentation step, replacing older workflows that required separate fragmentation and ligation reactions^15^. Because the Tn5 transposase is immobilized on streptavidin-coated magnetic beads at a fixed enzyme density, the reaction effectively saturates once all transposomes are occupied, providing a self-normalizing behavior over input DNA concentrations from low single-digit to high double-digit ng/µL and reducing the need for precise pre-tagmentation normalization^15^. Compared with earlier Nextera Flex (DNAprep) implementations that depended on tightly controlled input amounts, this bead-linked format stabilizes the enzyme on a solid support, improves handling and wash steps, and makes the workflow substantially more forgiving of variation in both DNA concentration and purity that is typical of wild plant collections^18^.

Following tagmentation, Tn5 remains bound to the adapter-tagged DNA, so incorporation of an SDS treatment at 55 °C denatures the enzyme and disrupt the protein-DNA complex, which both halts further transposition and releases fragments for efficient PCR amplification^19^. Subsequent PCR adds the full-length Illumina adapters and incorporates unique dual i5 and i7 index combinations, enabling multiplexing of up to 96 libraries per plate and allowing large sample sets to be pooled into single high-throughput sequencing runs while retaining sample identity through bioinformatic demultiplexing. In practice, this indexing strategy yielded consistently high read counts and low index mismatch rates across all eight validation libraries, with no obvious penalty for samples that began with lower A260/230 ratios, highlighting the protocol’s tolerance for imperfect DNA purity.

SPRI bead purification is used both to remove inhibitors carried over from extraction and to shape the final fragment size distribution via two-sided size selection. In the presence of polyethylene glycol(PEG) and NaCl, DNA becomes less soluble and binds the carboxylated paramagnetic beads in a size-dependent fashion, such that altering the bead-to-sample ratio selectively retains or excludes different fragment length classes^20^. By first applying a 0.5× SPRI ratio to remove large fragments and then a 0.75× ratio to capture fragments greater than approximately 300 bp, the pooled libraries were consistently enriched for inserts in the 300-700 bp range, with TapeStation analysis showing a primary peak near 371 bp and minimal off-target material, confirming that the empirically optimized ratios produce a tight, Illumina-compatible insert distribution.

Across eight wild and cultivated *Vitis* accessions, Nanodrop and Qubit measurements revealed broad variation in DNA yield and purity, with Qubit concentrations spanning roughly 2–60 ng/µL and several field samples exhibiting A260/230 values well below the theoretical clean DNA threshold of 2.0. Despite this heterogeneity, agarose gels showed predominantly high-molecular-weight genomic DNA after extraction, and all samples produced clear library smears prior to size selection and high-quality pooled libraries after SPRI cleanup, indicating that the combined CTAB-PTB extraction and bead-linked Tn5 library preparation effectively decoupled library success from strict absorbance-based purity metrics. Sequencing on a NovaSeq X platform further confirmed that read yield, index quality, and overall sequencing performance were not systematically compromised in samples with lower A260/230 ratios, supporting the conclusion that moderate levels of co-extracted metabolites are well tolerated by this workflow.

Because both the extraction reagents and in-house bead-bound Tn5 transposase are inexpensive^19^, the complete workflow can be executed in a 96-well plate format with a per-sample cost of roughly 3 USD and a turnaround time of approximately 2.5 days from tissue DNA extraction to pooled, size-selected libraries, making it readily scalable to hundreds of samples. This cost-effectiveness, combined with the ability to process DNA of variable quality without extensive normalization or repeated extractions, directly addresses logistical bottlenecks in population and landscape genomics, where discarding or re-extracting marginal samples can inflate costs and erode the environmental and geographic breadth of sampling. This protocol has been validated for DNA extracted from fresh wild grape (*Vitis* spp.) leaf tissue. Libraries were also successfully prepared for soybean samples; however, additional optimization may be needed for other plant taxa. Input DNA concentration must exceed 0.314 ng/ul, as quantified using a Qubit fluorometer. Each batch of in-house prepared Tn5 transposase and SPRI beads requires calibration before use to ensure consistent library quality. This method is not designed for highly degraded or herbarium DNA, which remains as an important boundary on its broader applicability.

Troubleshooting:

1. Low DNA yield: Reasons: not enough tissue taken, tissues not young enough, incomplete tissue homogenization by bead beater, not all tissues exposed to buffer for lysis during the mixing step, overdrying of DNA pellet during ethanol washes
2. Poor DNA quality: Reasons: impurities were also pipetted along with the DNA during the chloroform step, not enough ethanol washes
3. Fragmented DNA on gel Reasons: Shearing by excessive pipetting or vortexing
4. Library not visualized on gel: Tn5 did not come in contact with DNA, tn5 did not bind to beads, Tn5+beads lost function due to longer storage time, problems in amplification, and PCR steps.
5. Wrong library size: High tn5 activity means shorter fragments, so work on ice. SDS did not work due to precipitation
6. Wrong pooled library size: Wrong spri bead ratio- can do re-size selection, improper storage of spri beads leading to loss of function, pipetting error causing wrong volumes to be used.

## Key Resource table

**Table 13.** Key resources and their source used in the protocol.

| Reagent or Resource | Source | Identifier |
| --- | --- | --- |
| Cetyltrimethylammonium bromide 500 g (CTAB) | Fisher Scientific | MFCD00011772 |
| Bead Mill 24 Homogenizer | Fisher Scientific | 15-340-163 |
| Invitrogen™ PureLink™ RNase A (20 mg/mL) | Fisher Scientific/Invitrogen | 12091021 |
| Streptavidin beads | Fisher Scientific | S1421S |
| DMF (Dimethylformamide, 50ML) | Avantor Scientific | MFCD00003284 |
| Polymerase (Q5 High-Fidelity DNA Polymerase) | Avantor Scientific/New England Biolabs(NEB) | M0491L |
| dNTP (Deoxynucleotide (dNTP) solution mix- 8 umol) | NEB | N0447S |
| PTB(N-Phenacylthiazolium bromide 1g) | Oakwood Chemical | 080244-1g |
| 50 mL tube magnetic stand | Sergi Lab Supplies/Amazon | B08136XC2P |
| 0.2 % Triton X-100 | Fisher Scientific | MFCD00132505 |
| 200 ul Wide Bore tips | Fisher Scientific | 2707600 |
| Invitrogen™ Nuclease-Free Water | Fisher Scientific | AM9937 |
| EDTA 0.5 M pH 8 100 ml | Fisher Scientific | MFCD00003541 |
| 5 m Sodium Chloride (NaCl) 1 L | Fisher Scientific | MFCD00003477 |
| Tris-HCl 1 M pH 8 100 ml | Fisher Scientific | J22638-AE |
| Thermo Scientific™ TopVision Agarose 100g | Fisher Scientific | FERR0491 |
| Thermo Scientific™ PCR Plate, 96-well, semi-skirted, flat deck | Fisher Scientific | AB-1400 |
| Fisherbrand™ Colored Screw Caps for Fisherbrand™ Nonsterile Threaded End Microcentrifuge Tubes without Caps | Fisher Scientific | 2681364 |
| Fisherbrand™ Nonsterile Threaded End Microcentrifuge Tubes without Caps | Fisher Scientific | 2681344 |
| Thermo Scientific™ Adhesive PCR Plate Foils | Fisher Scientific | AB-0626 |
| Fisherbrand™ Mini Tube Rotator | Fisher Scientific | 88861051 |
| i5 oligos | Eurofins | Custom-made sequences |
| i7 oligos | Eurofins | Custom-made sequences |
| 96-well plate magnet | Sergi Lab Supplies/Amazon | B08134P9RT |
| Eppendorf™ ThermoMixer Temperature Control Device | Fisher Scientific/Eppendorf | 5382000023 |
| 2 ml magnetic rack | Sergi Lab Supplies/Amazon | B0CFXF5NKZ |
| Thermo Scientific™ GeneRuler DNA Ladder Mix | ThermoScientific Fermentas | FERSM0332 |
| Invitrogen™ SYBR™ Safe DNA Gel Stain | Invitrogen | S33102 |
| BioSpec Products 2.3 mm Zirconia/Silica Beads | Fisher Scientific/Biospec Products | 11079125Z |

Primers and Oligos:

**Table 14.**
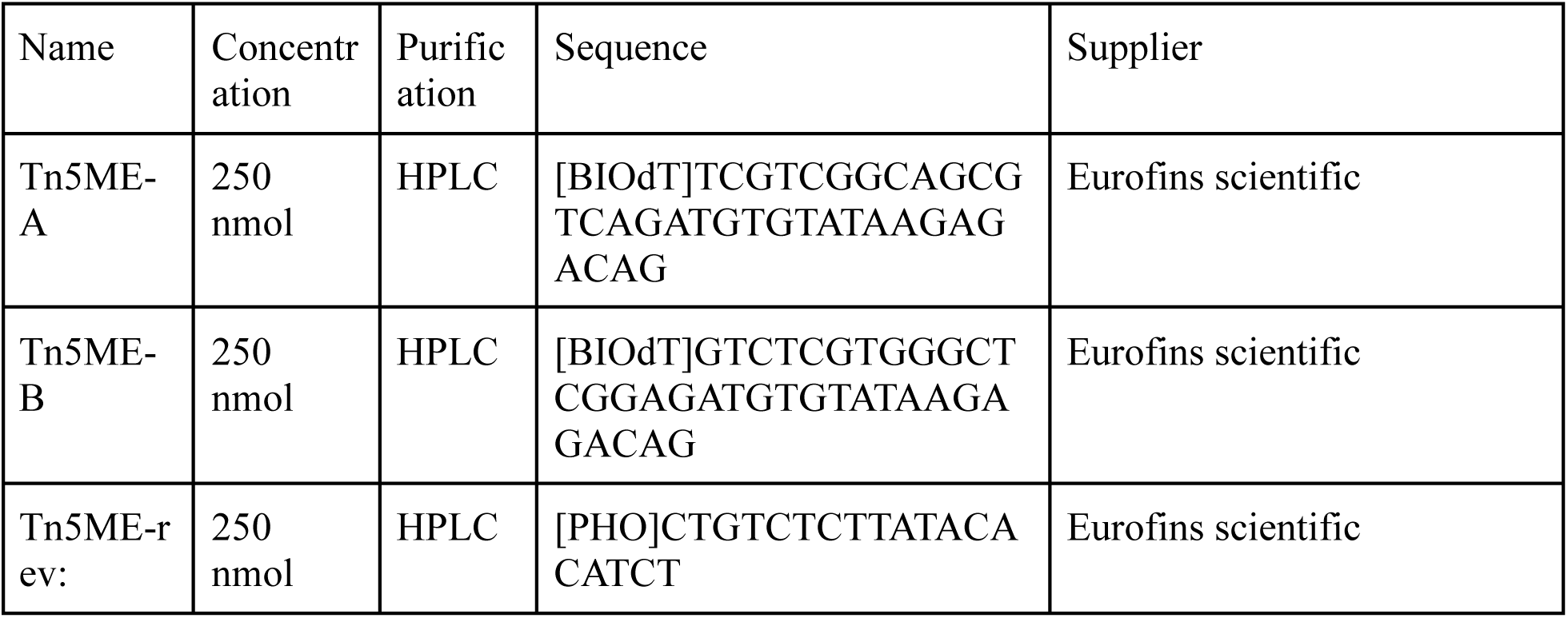
Biotinylated oligos used to bind to purified Tn5.

Resource Availability:

**Table 15.**
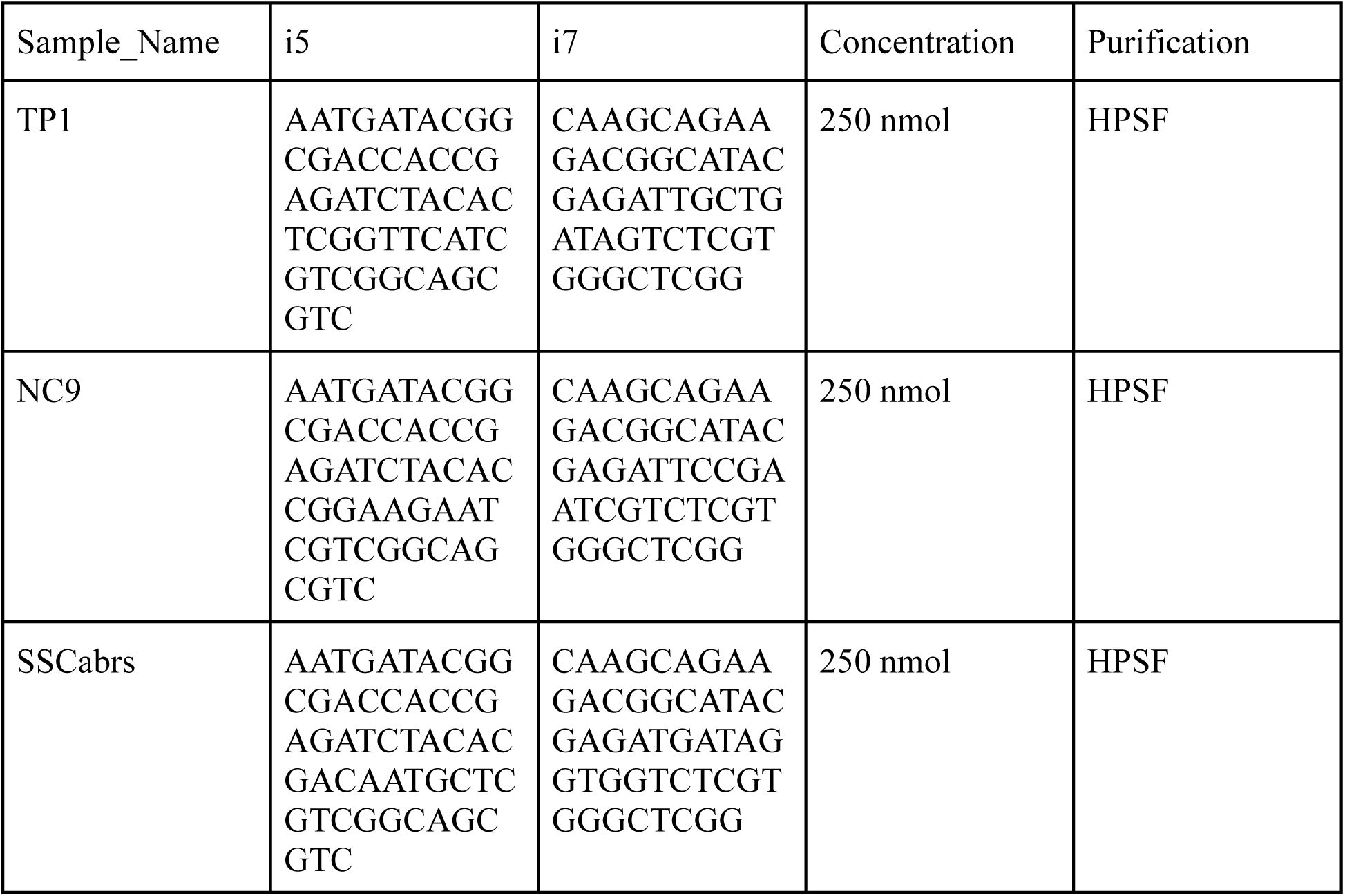
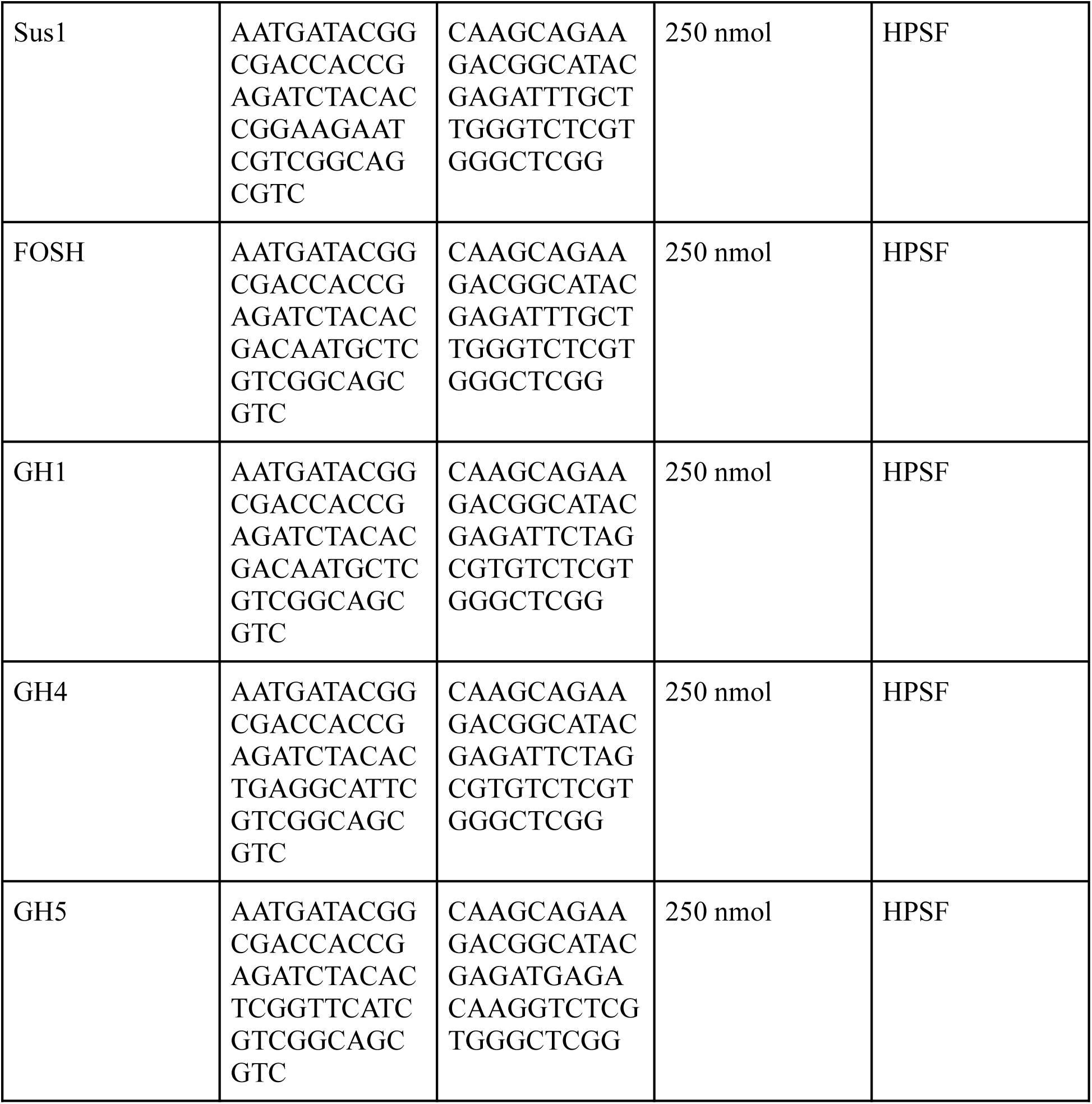
Unique dual indexes used for barcoding the DNA samples.

Data from the sequencing is available at https://github.com/gaushi/robust_vitis_wgs

## Notes

https://github.com/gaushi/robust_vitis_wgs

### Competing Interest Statement

The authors have declared no competing interest.

## References

1. Song, B.-H., and Mitchell-Olds, T. (2011). Evolutionary and ecological genomics of non-model plants. J. Syst. Evol. 49, 17–24. 10.1111/j.1759-6831.2010.00111.x.

2. Pasquali, F., Do Valle, I., Palma, F., Remondini, D., Manfreda, G., Castellani, G., Hendriksen, R.S., and De Cesare, A. (2019). Application of different DNA extraction procedures, library preparation protocols and sequencing platforms: impact on sequencing results. Heliyon 5, e02745. 10.1016/j.heliyon.2019.e02745.

3. Turner, S., Armstrong, L.L., Bradford, Y., Carlson, C.S., Crawford, D.C., Crenshaw, A.T., de Andrade, M., Doheny, K.F., Haines, J.L., Hayes, G., et al. (2011). Quality control procedures for genome-wide association studies. Curr. Protoc. Hum. Genet. Chapter 1, Unit1.19. 10.1002/0471142905.hg0119s68.

4. Santos, A.S., and Gaiotto, F.A. (2020). Knowledge status and sampling strategies to maximize cost-benefit ratio of studies in landscape genomics of wild plants. Sci. Rep. 10, 3706. 10.1038/s41598-020-60788-8.

5. Hübner, S., and Kantar, M.B. (2021). Tapping diversity from the wild: From sampling to implementation. Front. Plant Sci. 12, 626565. 10.3389/fpls.2021.626565.

6. Asik, E., Kashif, S.Z., and Oztas, O. (2025). Plant tolerance mechanisms to DNA-damaging UV stress. J. Exp. Bot. 77, 27–50. 10.1093/jxb/eraf272.

7. Schenk, J.J., Becklund, L.E., Carey, S.J., and Fabre, P.P. (2023). What is the “modified” CTAB protocol? Characterizing modifications to the CTAB DNA extraction protocol. Appl. Plant Sci. 11, e11517. 10.1002/aps3.11517.

8. Friar, E.A. (2005). Isolation of DNA from plants with large amounts of secondary metabolites. Methods Enzymol. 395, 3–14. 10.1016/S0076-6879(05)95001-5.

9. Zielińska, S., Radkowski, P., Blendowska, A., Ludwig-Gałęzowska, A., Łoś, J.M., and Łoś, M. (2017). The choice of the DNA extraction method may influence the outcome of the soil microbial community structure analysis. Microbiologyopen 6, e00453. 10.1002/mbo3.453.

10. Marsh, W.A., Hall, A., Barnes, I., and Price, B. (2025). Facilitating high throughput collections-based genomics: a comparison of DNA extraction and library building methods. Sci. Rep. 15, 6013. 10.1038/s41598-025-88443-0.

11. Rohland, N., and Reich, D. (2012). Cost-effective, high-throughput DNA sequencing libraries for multiplexed target capture. Genome Res. 22, 939–946. 10.1101/gr.128124.111.

12. Öncü Öner, T., Temel, M., Pamay, S., Abaci, A.K., and Kaya Akkale, H.B. (2023). An improved method for efficient DNA extraction from grapevine. International Journal of Life Sciences and Biotechnology 6, 21–36. 10.38001/ijlsb.1150387.

13. Doyle, J.J., and Doyle, J.L. (1987). A Rapid DNA Isolation Procedure for Small Quantities of Fresh Leaf Tissue. Phytochemical Bulletin 19, 11–15.

14. Marinček, P., Wagner, N.D., and Tomasello, S. (2022). Ancient DNA extraction methods for herbarium specimens: When is it worth the effort? Appl. Plant Sci. 10, e11477. 10.1002/aps3.11477.

15. Bruinsma, S., Burgess, J., Schlingman, D., Czyz, A., Morrell, N., Ballenger, C., Meinholz, H., Brady, L., Khanna, A., Freeberg, L., et al. (2018). Bead-linked transposomes enable a normalization-free workflow for NGS library preparation. BMC Genomics 19, 722. 10.1186/s12864-018-5096-9.

16. Kucka, M. (2020). Tn5-on-Beads DNA Tagmentation Protocol. >. https://policycommons.net/artifacts/1669150/tn5-on-beads-dna-tagmentation/2400799/.

17. Lucena-Aguilar, G., Sánchez-López, A.M., Barberán-Aceituno, C., Carrillo-Ávila, J.A., López-Guerrero, J.A., and Aguilar-Quesada, R. (2016). DNA source selection for downstream applications based on DNA quality indicators analysis. Biopreserv. Biobank. 14, 264–270. 10.1089/bio.2015.0064.

18. Illumina DNA Prep product line https://www.illumina.com/products/by-brand/dna-prep-portfolio.html.

19. Picelli, S., Björklund, A.K., Reinius, B., Sagasser, S., Winberg, G., and Sandberg, R. (2014). Tn5 transposase and tagmentation procedures for massively scaled sequencing projects. Genome Res. 24, 2033–2040. 10.1101/gr.177881.114.

20. SPRIselect Beads Size Selection https://www.beckman.com/reagents/genomic/cleanup-and-size-selection/size-selection/spriselect-protocol.

